# Active anaerobic methane oxidation and sulfur disproportionation in the deep terrestrial subsurface

**DOI:** 10.1101/2021.08.21.457207

**Authors:** Emma Bell, Tiina Lamminmäki, Johannes Alneberg, Chen Qian, Weili Xiong, Robert L. Hettich, Manon Frutschi, Rizlan Bernier-Latmani

## Abstract

Microbial life is widespread in the terrestrial subsurface and present down to several kilometers depth, but the energy sources that fuel metabolism in deep oligotrophic and anoxic environments remain unclear. In the deep crystalline bedrock of the Fennoscandian Shield at Olkiluoto, Finland, opposing gradients of abiotic methane and ancient seawater-derived sulfate create a terrestrial sulfate-methane transition zone (SMTZ). We used chemical and isotopic data coupled to genome-resolved metaproteogenomics to demonstrate active life and, for the first time, provide direct evidence of active anaerobic oxidation of methane (AOM) in a deep terrestrial bedrock. Proteins from *Methanoperedens* (formerly ANME-2d) are readily identifiable despite the low abundance (≤1%) of this genus and confirm the occurrence of AOM. This finding is supported by ^13^C-depleted dissolved inorganic carbon. Proteins from *Desulfocapsaceae* and *Desulfurivibrionaceae*, in addition to ^34^S-enriched sulfate, suggest that these organisms use inorganic sulfur compounds as both electron donor and acceptor. Zerovalent sulfur in the groundwater may derive from abiotic rock interactions, or from a non-obligate syntrophy with *Methanoperedens*, potentially linking methane and sulfur cycles in Olkiluoto groundwater. Finally, putative episymbionts from the candidate phyla radiation (CPR) and DPANN archaea represented a significant diversity in the groundwater (26/84 genomes) with roles in sulfur and carbon cycling. Our results highlight AOM and sulfur disproportionation as active metabolisms and show that methane and sulfur fuel microbial activity in the deep terrestrial subsurface.

**Significance Statement:** The deep terrestrial subsurface remains an environment in which there is limited understanding of the extant microbial metabolisms, despite its reported large contribution to the overall biomass on Earth. It is much less well studied than deep marine sediments. We show that microorganisms in the subsurface are active, and that methane and sulfur provide fuel in the oligotrophic and anoxic subsurface. We also uncover taxonomically and metabolically diverse ultra-small organisms that interact with larger host cells through surface attachment (episymbiosis). Methane and sulfur are commonly reported in terrestrial crystalline bedrock environments worldwide and the latter cover a significant proportion of the Earth’s surface. Thus, methane- and sulfur-dependent microbial metabolisms have the potential to be widespread in the terrestrial deep biosphere.

## Introduction

Microbial cells in the subsurface comprise a significant proportion of the global prokaryotic biomass (1). They are typically slow-growing, with long generation times ranging from months to years (2– 5). Despite their slow growth, they are active (6–9) and can respond to changes in subsurface conditions (10, 11). Understanding the drivers of microbial metabolism in the subsurface is therefore an important consideration for subsurface activities related to energy production, storage, and waste disposal.

The engineering of stable bedrock formations for industrial purposes (spent nuclear fuel disposal, underground storage of gases, geothermal energy production, oil and gas recovery) alters the subsurface ecosystem causing biogeochemical changes that impact microbial community activity and function (12, 13). Microbial activity can be beneficial, either through enhanced recovery (14), removal of unwanted products (15), or competitive exclusion (16). It is, however, often associated with detrimental impacts such as microbially-influenced corrosion (17), petroleum reservoir souring (18), or reduced permeability and clogging resulting from the formation of biofilms (13).

The crystalline bedrock of Olkiluoto, Finland, will host a repository for the final disposal of spent nuclear fuel. Copper canisters containing spent nuclear fuel will be one of several engineered barriers for safe final disposal, the final barrier being the Olkiluoto bedrock (400–500 m depth). Understanding the energy sources that fuel microbial activity at this site is important as slowly moving groundwater present in cracks and fractures in the bedrock has the potential to reach and interact with the engineered barriers. The microbial transformation of sulfate to sulfide in the fracture water is a significant concern as it could accelerate corrosion of copper canisters and compromise their longevity. The groundwater at Olkiluoto is geochemically stratified with depth with a salinity gradient that extends from fresh near the surface (<100 m) to saline (>1000 m). Sulfate is found in fractures at ∼100–300 m depth. Below 300 m, the concentration of sulfate declines in an opposing trend to methane, which increases with depth. This creates a sulfate-methane transition zone (SMTZ) at ∼250–350 m depth.

Sulfide is detected in relatively few drillholes at Olkiluoto, and its presence tends to be in groundwater from fractures in the SMTZ (19). Exceptions in deeper fractures exist where drilling has resulted in the introduction of sulfate-rich groundwater to deep methane-rich groundwater providing an electron acceptor to a system otherwise limited in terminal electron acceptors (11). In marine and coastal sediments, SMTZs are host to anaerobic methanotrophic archaea (ANME) and sulfate-reducing bacteria that syntrophically cycle methane and sulfate (20), and quantitatively consume methane diffusing up from deep methanogenic sources through anaerobic oxidation of methane (AOM) (21, 22). In contrast, the biogeochemical significance of syntrophic interactions is poorly understood in terrestrial SMTZs. Previous work has proposed, but not demonstrated, widespread AOM in crystalline bedrock environments. This hypothesis was based on the detection of ANME from the archaeal family *Methanoperedenaceae* (formerly ANME-2d) in groundwater from granitic bedrock in both Finland (23, 24) and Japan (25–27). Additionally, past AOM activity has also been evidenced by the precipitation of extremely ^13^C-depleted calcites in the granitic bedrock of Sweden (28).

While many ANME rely on a syntrophic partner to couple methane oxidation to the reduction sulfate, *Methanoperedenaceae* can independently conduct methane oxidation with diverse terminal electron acceptors (29, 30). *Methanoperedenaceae* were originally enriched in a bioreactor shown to couple AOM to the reduction of nitrate (31). *Methanoperedenaceae* lacking nitrate reductase genes have since been enriched in bioreactors demonstrated to couple AOM to the reduction of iron (Fe(III)) and manganese (Mn(IV)) (32). Studies from freshwater lake sediments have proposed that *Methanoperedenaceae* also participate in sulfate-dependent AOM (33, 34). Similarly, enrichment experiments using deep groundwater suggest *Methanoperedenaceae* conduct sulfate-dependent AOM in the terrestrial subsurface (26, 35). However, AOM activity has not been demonstrated *in situ* in any deep crystalline bedrock terrestrial environment to date.

Here, we use genome-resolved metaproteogenomics to uncover active sulfur and methane cycling microorganisms in groundwater from a fracture in the SMTZ at Olkiluoto, Finland. Our goal was to constrain the electron donor(s) fueling sulfidogenesis and determine whether sulfur and methane cycles in the terrestrial SMTZ are linked. Stable isotopes and metaproteomics show that active AOM occurs in this environment and that sulfate-reducing bacteria (SRB) and sulfur-disproportionating bacteria (SDB) are active. Our data suggest that a syntrophic relationship may exist between ANME archaea and *Desulfobacterota* with dissimilatory sulfur metabolism, either through extracellular electron transfer or diffusible sulfur species. Additionally, we propose that ultra-small bacteria and archaea, abundant in the groundwater, live as episymbionts on the primary microbiome.

## Results

### Stable isotopes indicate occurrence of dissimilatory sulfur metabolism and anaerobic oxidation of methane

The groundwater was temperate (9.6°C), reducing (oxidation-reduction potential = −420 mV), brackish (total dissolved solids (TDS) 7.4 g/L; conductivity 13 mS/cm) and at circumneutral pH (pH value 7.6). DAPI (4′6-diamidino-2-phenylindole) cell counts indicated an abundance of 2.3 × 10^5^ cells/mL. Sulfate in the groundwater (∼0.8 mM; Fig. 1A) originates from the Littorina Sea, which preceded the Baltic Sea 2,500–8,500 years ago (19). In the sampled fracture, sulfate is ^32^S-depleted (44.6 ‰ VCDT; Fig. 1A) relative to groundwater with the same sulfate source at Olkiluoto (∼25 ‰; (19)) resulting from the preferential use of ^32^S by microorganisms with dissimilatory sulfur metabolism. Reduced sulfur compounds, thiosulfate (∼0.1 mM) and sulfide (∼0.4 mM) are also present (Fig. 1A), indicating ongoing metabolism of sulfur compounds. The isotope composition of methane (−42.9 ‰ VPBD; Fig. 1b) reflects deep abiotic methane mixed with an increasing fraction of microbially-produced methane with decreasing depth at Olkiluoto (19). During AOM, the light carbon isotope value of methane is transferred to CO_2_. The light dissolved inorganic carbon (DIC) isotope value (−28.0 ‰ VPBD; Fig. 1B) therefore indicates that methane could have contributed to the DIC pool, and likely reflects the mixing of distinct carbon sources including a minor AOM part. Hydrogen was relatively abundant (∼30 µM) throughout the sampling period, except in the final sampling month (November) where hydrogen was not detected (Dataset S1). The concentration of dissolved gases *in situ* may be greater than the measured values as the pressure decrease during pumping can cause degassing. Acetone (6 µM) and ethanol (3 µM) were detected but account for only ∼5% of the measured dissolved organic carbon (0.5 mM), meaning that unidentified organic compounds are present in the groundwater.

**Figure 1.**
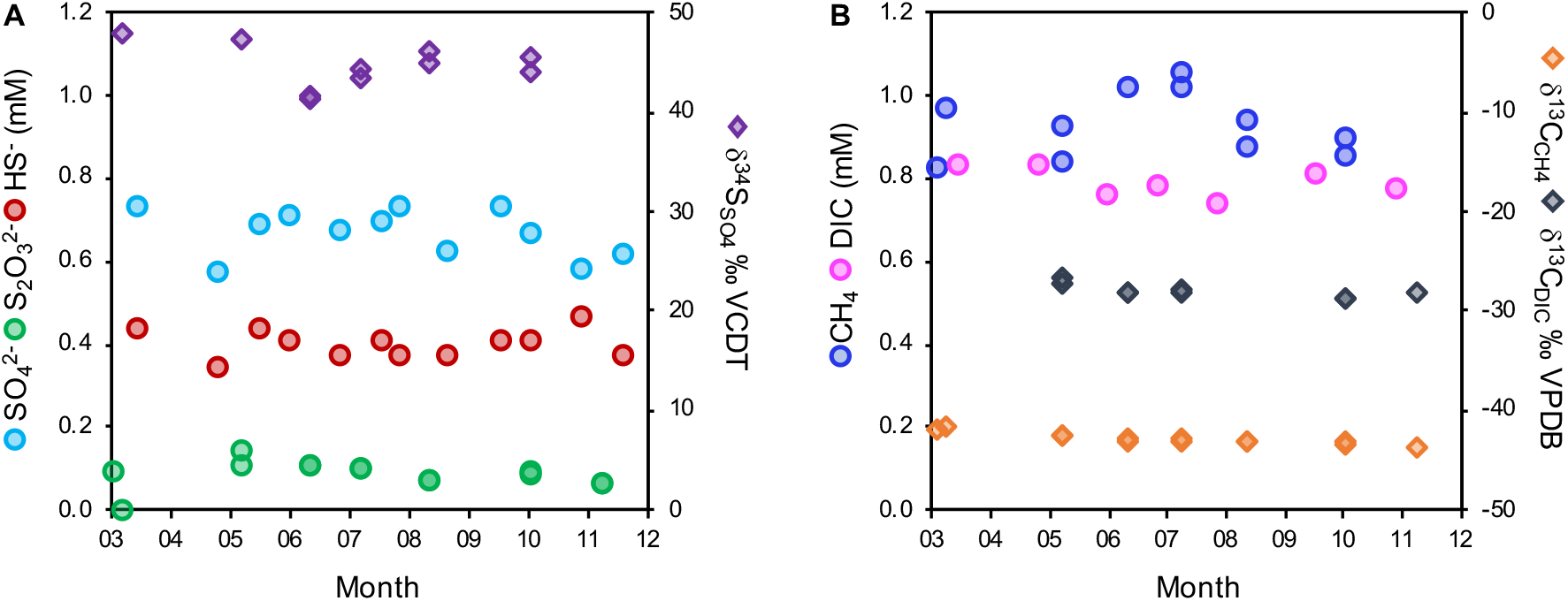
Groundwater chemistry (drillhole OL-KR13). (A) Sulfur species and sulfur isotope values (^34^S/^32^S expressed as δ^34^S_SO4_); (B) dissolved inorganic carbon (DIC), total organic carbon (DOC) and methane. Carbon isotope values (^13^C/^12^C) are expressed as δ^13^C_DIC_ and δ^13^C_CH4_. Data used in this figure is provided in Dataset S1.

### Metaproteomics reveal active sulfur disproportionation among sulfur cycling bacteria

Analysis of the microbial community composition by 16S rRNA gene amplicon profiling (Fig. 2) and phylogenomics (Fig. 3) revealed diverse microorganisms from both bacterial and archaeal lineages. Dereplication of metagenome assembled genomes (MAGs) from seven metagenomic datasets augmented with single cell genomes (SAGs) resulted in the recovery of 84 genomes from 25 phyla (Fig. 3; genome completeness is provided in Dataset S2). *Desulfobacterota*, a phylum known to harbor sulfur-cycling bacteria, were most abundant in the 0.2 µm size fraction (Fig. 2), consistent with chemical and isotope data that support dissimilatory sulfur metabolism in the groundwater (Fig. 1). Genes for dissimilatory sulfate reduction; sulfate adenylyltransferase (*sat*), adenylylsulfate reductase alpha and beta subunits (*aprAB*), and dissimilatory sulfite reductase alpha, beta and delta subunits (*dsrABD*), were detected in MAGs from the phyla *Desulfobacterota* and *Nitrospirota* (Fig. 4). Genomes from these phyla also contained electron transport complexes *qmoABC* and *dsrMKJOP* and the sulfur relay protein *dsrC* (Fig. 4 and Dataset S3). One MAG from the *Gammaproteobacteria* (genus *Hydrogenophaga*) contained genes for dissimilatory sulfate reduction (Dataset S3) but was inferred to be a sulfur oxidizer based on the presence of subunits *dsrEFH* (essential for reverse function) and absence of *dsrD* (needed for forward function) (36, 37).

**Figure 2.**
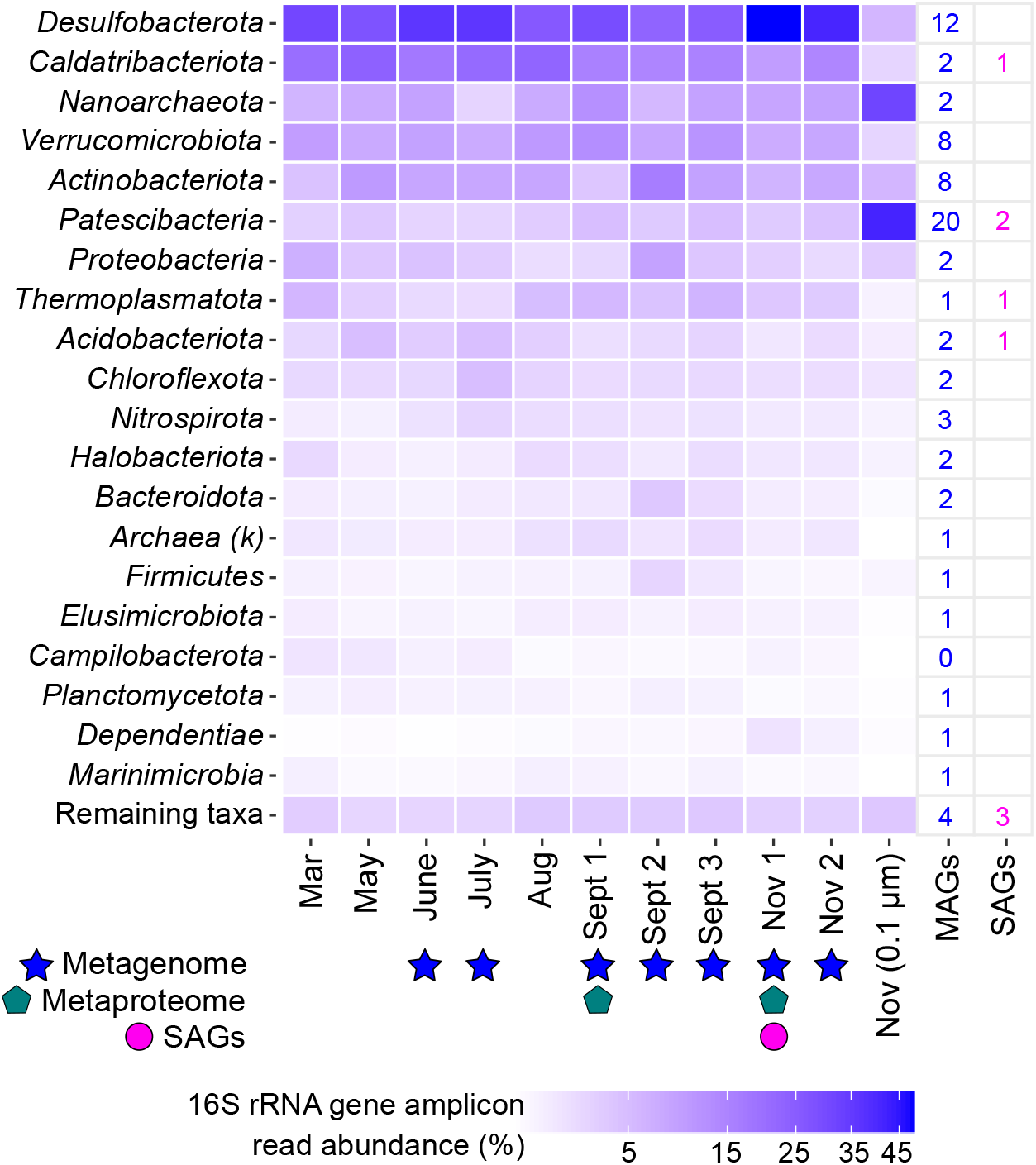
Phyla detected by 16S rRNA gene amplicon sequencing of groundwater from drillhole OL-KR13. DNA was extracted from biomass collected on an 0.2 µm filter for all samples except one (Nov (0.1 µm)). If the phylum could not be assigned, the kingdom is given, denoted by a ‘k’ in parentheses. Sampling months (March – November) with a corresponding metagenome (filled star), metaproteome (filled pentagon) and SAGs (filled circle) are indicated. The number of MAGs and SAGs is given for each phylum following dereplication. *Verrucomicrobiota* includes MAGs from the class *Omnitrophia* in the Silva 138 release that are included as a separate phylum (*Omnitrophota*) in the GTDB release 95. Genomes assigned to remaining taxa are: *Altiarchaeota* (MAG + SAG); *Bacteria UPB18* (2x MAGs); *Delongbacteria* (MAG); *Bipolaricaulota* (SAG); *Cloacimonadota* (SAG).

**Figure 3.**
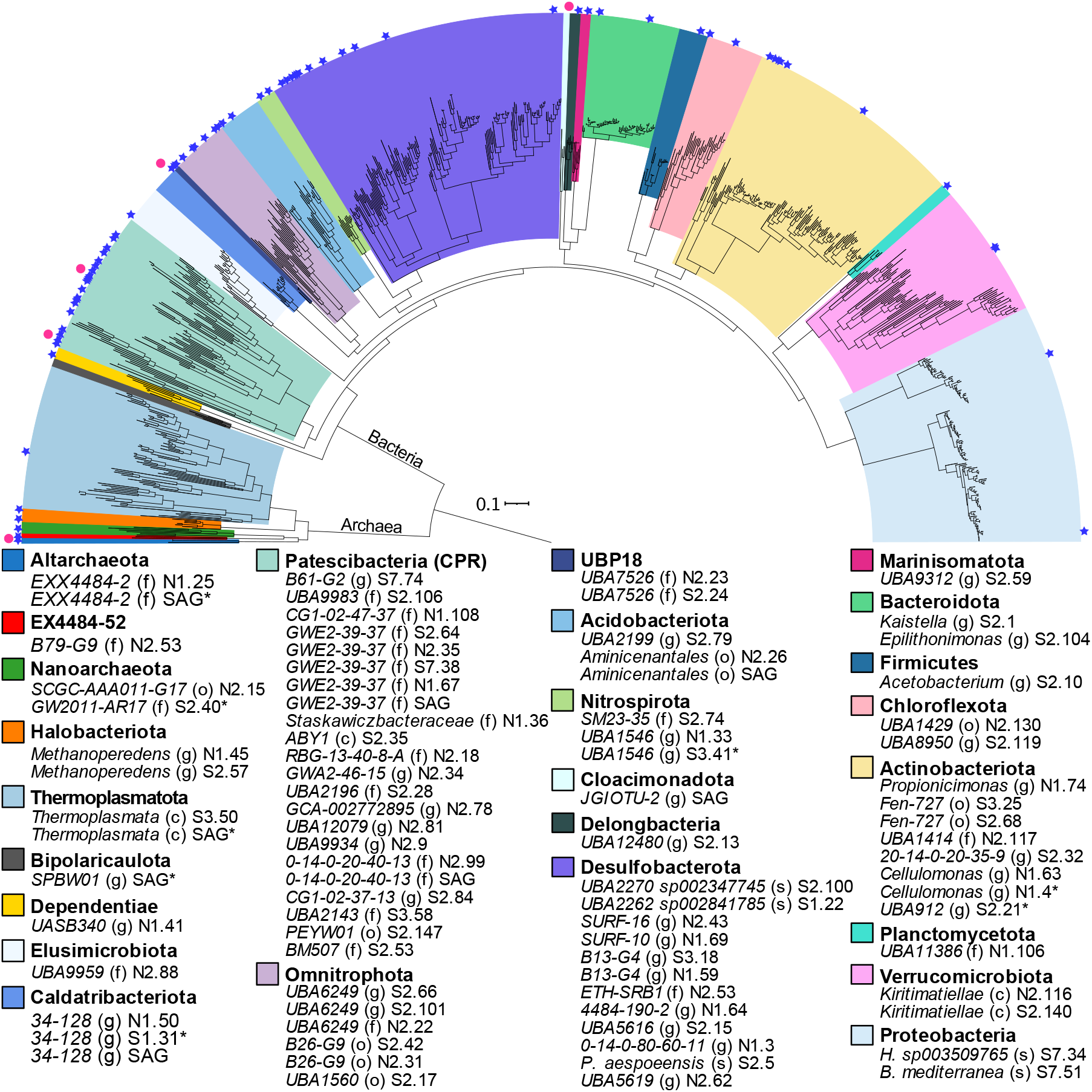
Diversity of recovered genomes (dereplicated MAGs and SAGs) based on 16 single copy genes. MAGs are indicated by a blue star and SAGs are indicated by a pink circle. The colors represent different phylum-level lineages. Letters in parentheses indicate the taxonomic rank assignment: o, order; c, class; f, family; g, genus; s, species. Full taxonomic assignments are provided in Dataset S2. MAGs/SAGs marked with an asterisk (*) were excluded from the tree as they contained too few of the single copy genes used for alignment. The scale bar corresponds to per cent average amino acid substitution over the alignment.

**Figure 4.**
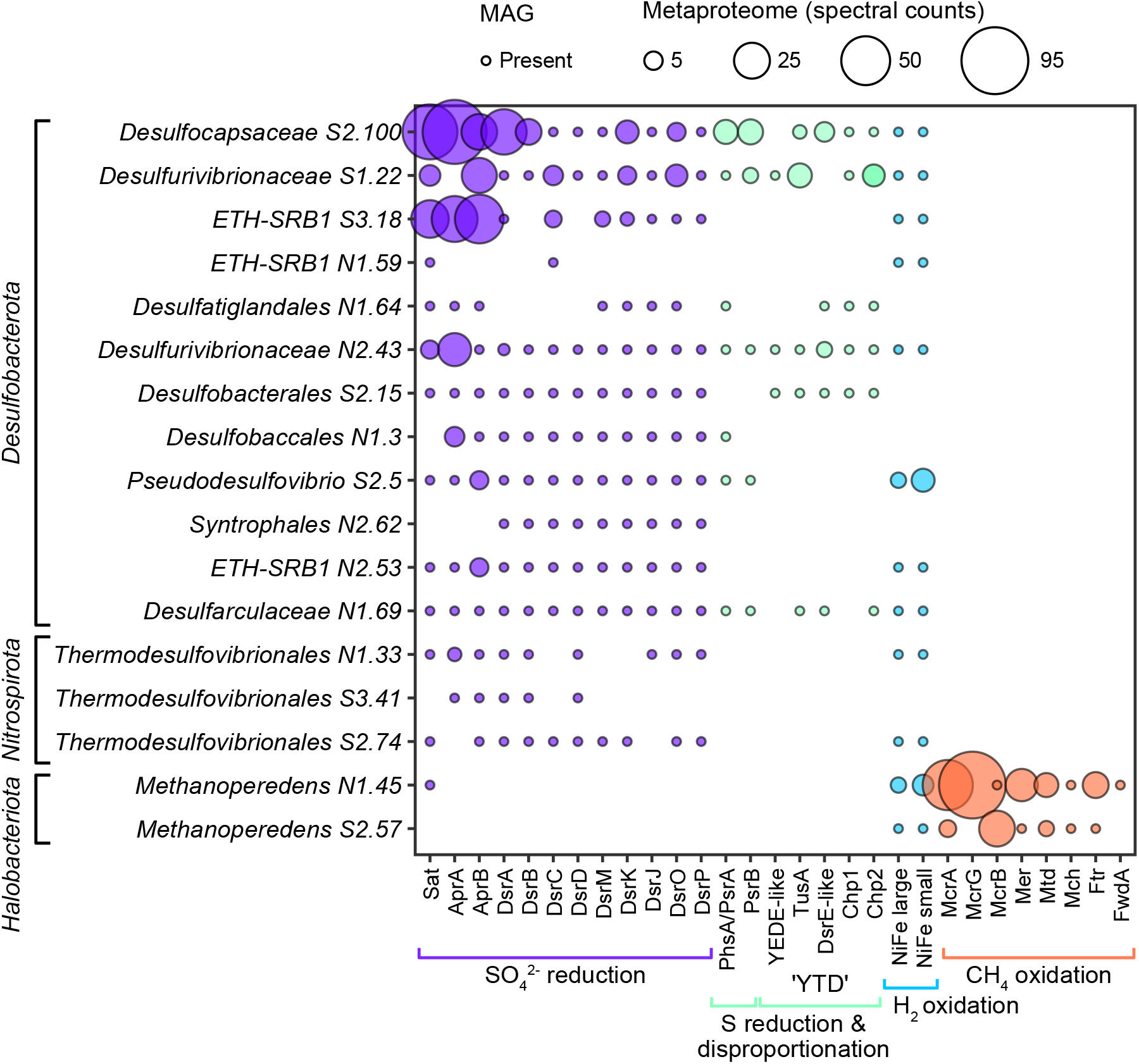
Abundance of proteins involved in sulfur, hydrogen and methane metabolism in *Desulfobacterota, Nitrospirota* and *Halobacteriota*. Filled circles indicate gene presence in the MAG. The size of the circle shows the abundance of the protein (spectral counts) in the metaproteome.

Metaproteomes from two sampling months (Fig. 4) were explored to uncover the activity of microorganisms in the groundwater at the time of sampling. The greatest number of proteins among sulfur-cycling bacteria were recovered from *Desulfocapsaceae, Desulfovibrionaceae*, and *ETH SRB-1* (Dataset S4). Sulfur cycling enzymes from these three MAGs had high spectral counts (the sum of MS/MS spectra for all peptides identified to a particular protein) indicating that these proteins were abundant (Fig. 4). *Desulfocapsaceae, Desulfovibrionaceae*, and *ETH SRB-1* and many other *Desulfobacterota* harbored the metabolic potential for H_2_ oxidation, but peptides from respiratory NiFe hydrogenase were only detected from *Pseudodesulfovibrio aespoeensis* (Fig. 4). *P. aespoeensis* was originally isolated from another subsurface location in the Fennoscandian Shield (Äspö, Sweden) (38), and belongs to the core microbiome of the aquifer in this crystalline rock (9). Peptides from the beta subunit of adenylylsulfate reductase (AprB) were also detected in the *P*.*aespoeensis* proteome (Fig. 4), thus, H_2_-dependent sulfate reduction is suggested as the metabolism for this MAG.

In addition to hydrogen, some *Desulfobacterota* had the potential to oxidize organic carbon (lactate and acetate; Dataset S3) but peptides from lactate dehydrogenase (Ldh) and acetyl-CoA synthetase (Acs) were not detected in the metaproteome. We therefore searched the metaproteome for enzymes catalyzing the oxidation of an alternate electron donor. We found that some proteins from the ‘YTD’ gene cluster were expressed in both *Desulfocapsaceae* and *Desulfovibrionaceae* (Fig. 4). This gene cluster is composed of five genes: a sulfur transport *yedeE*-like gene, a sulfurtransferase *tusA* gene, a *dsrE-*like gene, and two conserved hypothetical proteins (*chp1* and *chp2*) and is a potential genome marker for SDB (39). While it is difficult to discriminate SDB from SRB by genomic features alone (39–43), the expression of proteins from the YTD cluster suggest that *Desulfocapsaceae* and *Desulfovibrionaceae* (and possibly other *Desulfobacterota* with the YTD cluster; Fig. 4) disproportionate sulfur in this groundwater. *Desulfocapsaceae* and *Desulfovibrionaceae* also expressed a polysulfide reductase subunit B (*psrB*) adjacent to a thiosulfate reductase/polysulfide reductase (*phsA/psrA*) alpha subunit (Fig. 4). Polysulfide reductase catalyzes the respiratory conversion of polysulfide to hydrogen sulfide (44), while thiosulfate reductase catalyzes the initial step in disproportionation of thiosulfate (40). These enzymes are difficult to distinguish by their protein sequence as both are molybdopterin enzymes with a close phylogenetic relationship (45), but suggest that *Desulfocapsaceae* and *Desulfovibrionaceae* can use polysulfide and/or thiosulfate.

### Metaproteomics reveals active AOM in deep groundwater

*Methanoperedens* (phylum *Halobacteriota*) accounted for just ∼1% of 16S rRNA gene amplicon reads (Fig. 2) and the metagenomic community but had one of the greatest number of proteins detected in the metaproteome of all the recovered MAGs (Dataset S2). Proteins for AOM were abundant (Fig. 4), suggesting active AOM by *Methanoperedens* in this groundwater. Peptides from alpha and gamma subunits of methyl coenzyme M reductase (MCR) were also detected from a second, less abundant *Methanoperedens* MAG (0.17% of the metagenomic community). Furthermore, a contribution of methane-derived CO_2_ from *Methanoperedens* archaea is supported by the light δ^13^C_DIC_ signature (Fig. 1).

*Methanoperedens* had the metabolic potential for extracellular electron transfer, which could be either to an external electron acceptor or to a syntrophic partner. The potential for metal-dependent AOM was predicted by the presence of gene clusters encoding a NrfD-like transmembrane protein, a 4Fe-4S ferredoxin iron-sulfur protein, and multiheme cytochromes (MHC) (Dataset S5), hypothesized to mediate electron transfer from the cytoplasm to the periplasm in *Methanoperendaceae* during AOM coupled to Fe(III) and Mn(IV) oxides (29, 32). The more abundant *Methanoperedens* MAG also contained two MHC annotated as OmcX, an outer membrane protein necessary for electron transport out of the cell and growth on extracellular electron acceptors in *Geobacter* species (46). One copy of OmcX was located near an MHC with 28 heme-binding motifs predicted to be located extracellularly. Extracellular MHC could facilitate the transfer of electrons to metal oxides or to a syntrophic partner (47, 48). Archaeal flagellin have also been implicated in extracellular electron transfer by ANME archaea (49). Peptides from the major subunit of archaeal flagellin (FlaB) were detected in the metaproteome and may facilitate extracellular electron transfer to a syntrophic partner. Nitrate-dependent AOM was ruled out in this groundwater as nitrate reductase genes from *Methanoperedens* were absent in the two MAGs (Dataset S3). There was also no evidence of genes for the reduction of arsenate, selenate or elemental sulfur, found in some organisms from the *Methanoperedenaceae* family (29).

*Methanoperedens* encoded a respiratory NiFe group 1a hydrogenase, and peptides from both the large and small subunit were detected in the metaproteome (Fig. 4). The gene arrangement was the same as a *Methanoperedenaceae* MAG recovered from groundwater in Japan (26), where the NiFe 1a catalytic subunit is adjacent a *b*-type cytochrome and a hydrogenase maturation protease. In addition to the group 1 respiratory hydrogenase, *Methanoperedens* encoded a NiFe group 3b hydrogenase (Dataset S5), which is predicted to oxidize NADPH and evolve hydrogen (50). Some complexes have also been shown to have sulfhydrogenase activity, whereby the HydDA subunits encode a hydrogenase and the HydBG subunits encode the sulfur reductase component (51, 52).

### Candidate phyla radiation (CPR) bacteria and DPANN archaea contribute to carbon and sulfur transformations

Organisms from the superphyla *Patescibacteria* (also the CPR) and DPANN, which are known to have ultra-small cell sizes (53, 54), dominated the 0.1 µm size fraction (Fig. 2). Accordingly, genomes recovered from organisms belonging to CPR and DPANN organisms ranged from 0.6–1.7 Mbp (Dataset S2). Limited metabolic capabilities were evident (Fig. 5), consistent with the predicted symbiotic lifestyle of these lineages with other, larger microorganisms (55–57). In total, 87 proteins were detected in the metaproteome from CPR and DPANN organisms (Dataset S6). Most (65/87) were hypothetical or uncharacterized proteins, so putative function was predicted based on conserved domains in the protein sequence and/or sequence similarity (Dataset S6). The most abundant CPR proteins shared sequence similarity to protein sequences from other CPR organisms annotated as cell wall surface anchor family proteins. These abundant but uncharacterized proteins may therefore reflect the putatively episymbiotic lifestyle of CPR organisms and facilitate attachment to the host. Other abundant proteins had peptidoglycan-binding protein domains which are found in enzymes involved in bacterial cell wall degradation and phage endolysins (58). Type 4 pilus subunits (PilA and PilO) were also detected, which can enable attachment to surfaces, as well as potential host cells (59). Proteins recovered from CPR and DPANN organisms were also involved in cell core machinery and genetic information processing, such as those encoding ribosomal proteins, RNA-binding proteins, molecular chaperones, and elongation factors (Dataset S6).

**Figure 5.**
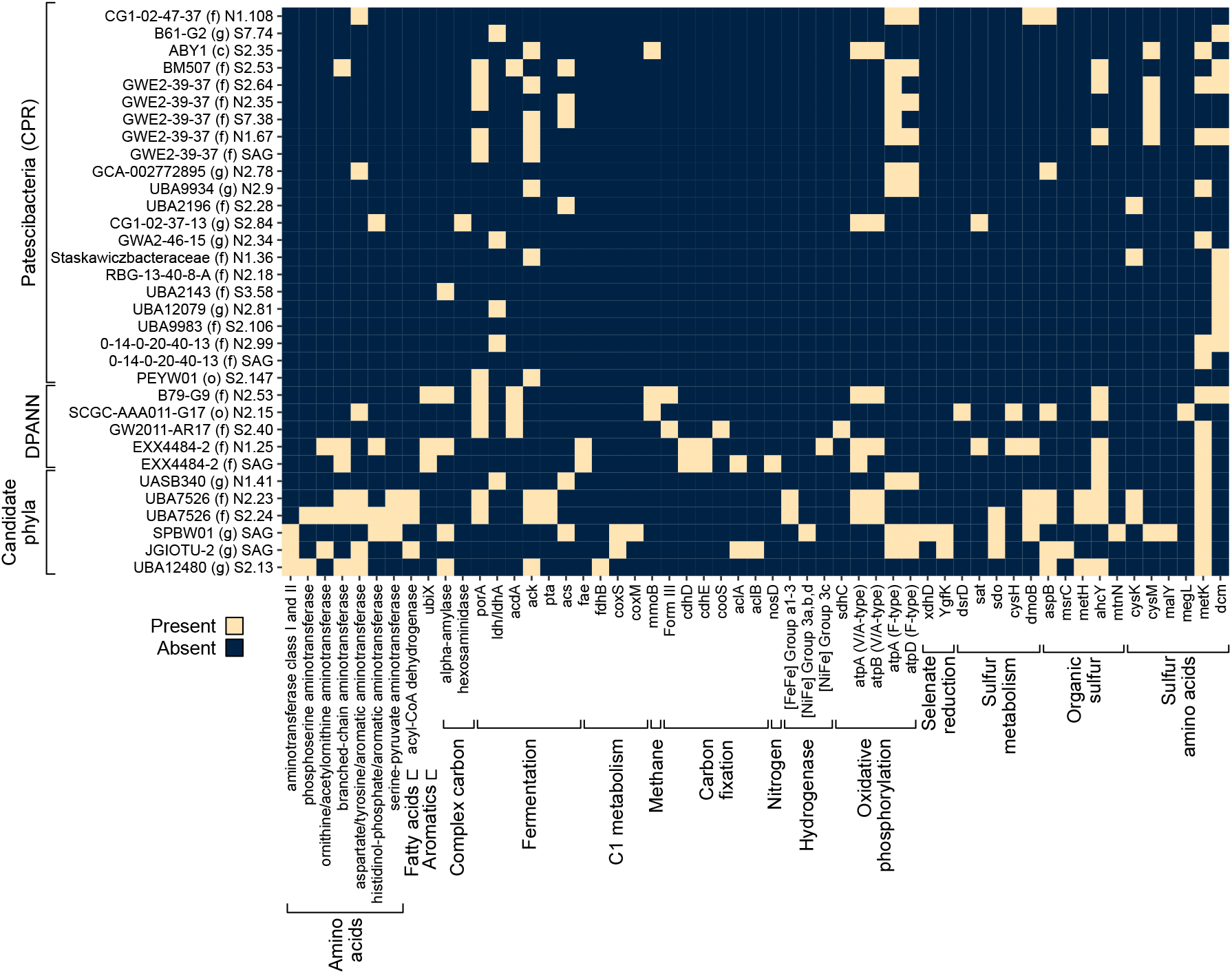
Metabolic profile of ultra-small CPR bacteria, DPANN archaea and other candidate phyla bacteria. Heatmap (presence/absence) of metabolic genes in MAGs and SAGs in Olkiluoto groundwater. Full gene names and enzyme reactions are provided in Dataset S3.

*Candidatus* Shapirobacterales BG1-G2, a member of the CPR supergroup *Microgenomates*, was one of the more abundant CPR detected, representing 1.8% of the microbial community (Dataset S2). *Cand*. Shapirobacterales expressed proteins with Type I and II cohesion domains in addition to a Type III dockerin repeat domain (Dataset S6), required for the formation of a cellulosome (a cellulose-degrading complex), indicating that this organism is actively involved in the metabolism of organic carbon. Organic carbon metabolism by this organism is further supported by the presence of glycoside hydrolases and polysaccharide lyases in the *Cand*. Shapirobacterales genome (Dataset S7). *Cand*. Shapirobacterales encoded lactate dehydrogenase (Fig. 5), indicating organic carbon could be fermented to lactate, as reported for other members of the CPR (60).

Other CPR and DPANN organisms also had the metabolic potential to supply fermentation products to their host cells. Fermentative metabolism (pyruvate oxidation and acetogenesis) was most common among the GWE2-39-37 family of *Patescibacteria* and was also detected in DPANN genomes (Fig. 5). DPANN archaea had genes for carbon fixation, including RuBisCo Form III (Fig. 5), which is predicted to function in the adenosine monophosphate pathway allowing DPANN to derive energy from ribose produced by other community members (61). *Patescibacteria* and DPANN organisms encoded sulfate adenylyltransferase (*sat*), dissimilatory sulfite reductase delta subunit (*dsrD*), phosphoadenosine phosphosulfate reductase (*cysH*), cysteine synthase (*cysK*), and S-sulfo-L-cysteine synthase (*cysM*), suggesting a potential role in sulfur cycling and the metabolism of sulfur amino acids (cysteine and methionine) as well as other organic sulfur compounds (Fig. 5). The metabolic capacity of CPR and DPANN symbionts has been shown to vary according to the metabolism of their host organisms (62). Thus, the potential for sulfur transformations by some CPR and DPANN could suggest a *Desulfobacterota* or *Nitrospirota* host in this groundwater.

## Discussion

Metaproteogenomic data reveal active sulfur and methane cycling microorganisms in deep groundwater in a terrestrial SMTZ. Canonical sulfate reducers coupling the oxidation of hydrogen to the reduction of sulfate are active, but metaproteomic data indicate that the most abundant *Desulfobacterota* in the groundwater gain energy from the disproportionation of inorganic sulfur (elemental sulfur or polysulfide). These sulfur compounds could be supplied by abiotic rock interactions (Fig. 6). For instance, exposure of iron silicate minerals from Olkiluoto bedrock to sulfide has been shown to result in the production of elemental sulfur and Fe^2+^ (which precipitates with excess sulfide), due to the abiotic reduction of Fe(III) in the major iron-bearing bedrock minerals (biotite, garnet and chlorite) (63). Inorganic sulfur could also be supplied through a syntrophic relationship with *Methanoperedens* in the form of diffusible sulfur species (Fig. 6), as proposed for other ANME archaea (64). In that case, both methane oxidation and sulfate reduction to zerovalent sulfur is carried out by *Methanoperedens*. The produced zerovalent sulfur (elemental sulfur and polysulfide) then provides both electron donor and acceptor to SDB, producing sulfate and sulfide (Fig. 6). Close physical association would not be required for syntrophy via diffusible sulfur species, suggesting that the bacterial partner may be non-specific. Alternatively, syntrophy between *Methanoperedens* and *Desulfobacterota* may be through extracellular electron transfer to SRB. Three MAGs belonging to the lineage ETH-SRB1 were recovered (Fig. 3), and sulfate reducers from this family have been suggested to form a syntrophic relationship with ethane-oxidizing archaea also from ANME-2d (65). ETH-SRB1 expressed enzymes for sulfate reduction but no enzymes for oxidation of an electron donor were detected in the metaproteome. Thus, ETH-SRB1 could be possible partners for *Methanoperedens* in this groundwater (Fig. 6).

**Figure 6.**
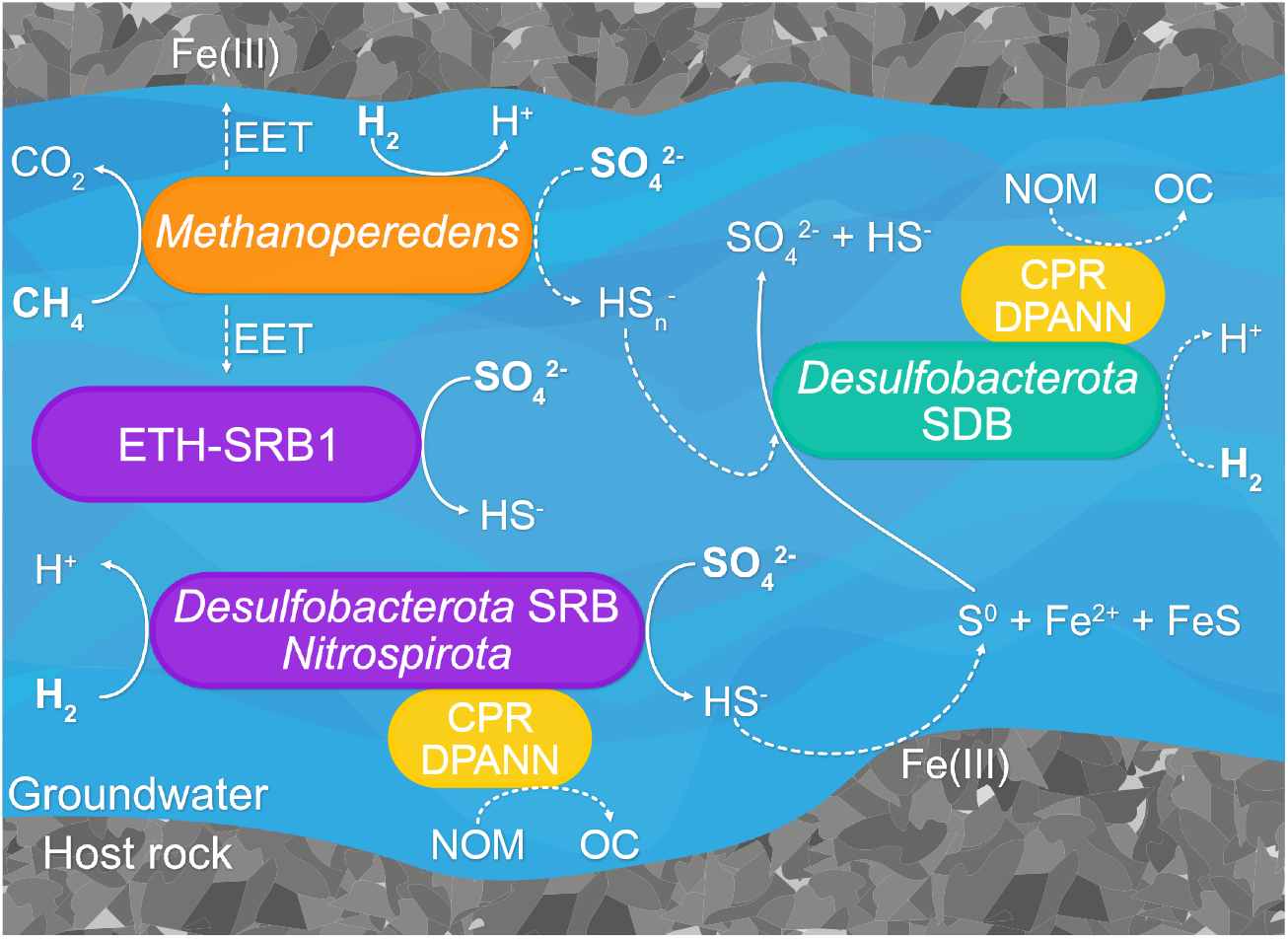
Proposed sulfur and methane cycling in Olkiluoto groundwater. Solid lines indicate metaproteomic evidence for the processes depicted while dashed lines indicate putative processes consistent with the data. Bold text indicates substrates present in the groundwater. Abbreviations shown in the figure are as follows: SRB, sulfate-reducing bacteria; SDB, sulfur-disproportionating bacteria; EET, extracellular electron transfer; NOM, natural organic matter; OC, organic carbon.

The genomic potential to transfer electrons extracellularly suggests that metal-dependent AOM could represent another metabolic strategy for *Methanoperedens* in Olkiluoto groundwater. Iron-bearing minerals in the bedrock have been shown to support culturable metal-reducing bacteria from Olkiluoto (63), and other metal-reducing bacteria have been shown to be active *in situ* (7). *Methanoperedens* expressed a respiratory hydrogenase suggesting active hydrogen oxidation. The potential for hydrogen oxidation by *Methanoperedenaceae* has been detected in other MAGs recovered from the deep terrestrial subsurface (26, 27), but this metabolic trait is not common amongst all genome-sequenced *Methanoperedenaceae* (29). The presence of a respiratory hydrogenase in some *Methanoperedenaceae* led to the suggestion that this metabolism may be advantageous in environments with temporally variable availability of methane (29). In this groundwater, however, *Methanoperedens* expresses both the core AOM pathway and the respiratory hydrogenase when both methane and hydrogen are available. The expression of both the core AOM pathway and the respiratory hydrogenase raises the possibility of hydrogenotrophic methanogenesis as a possible metabolism for this *Methanoperedens*, although this organism may simply gain energy from both methane and hydrogen. To our knowledge, *Methanoperedenaceae* have not been shown to produce methane, however, it has recently been suggested that ANME-1 archaea could alternate between AOM and methanogenesis, based on their presence in methane-producing sediments where hydrogen accumulates sufficiently to make hydrogenotrophic methanogenesis exergonic (66). A small shift in ^13^C_CH4_ toward lighter values was observed over the 12-month sampling period (Fig. 1), which indicates an increased proportion of ^12^C, associated with a biogenic methane contribution. The only organisms containing McrA and the capacity to produce methane in the metagenome were *Methanoperedens*. This methane isotope trend could, however, also be the result of long-term pumping of the fracture which also resulted in a shift in the groundwater chemistry overtime (Dataset S1). Furthermore, the light isotope ratio of dissolved inorganic carbon supports the contribution of methane-derived carbon. In ANME-1 archaea that are predicted to produce methane, it is not known whether one cell alternates between AOM and methanogenesis, or if different subclades selectively perform AOM or methanogenesis (66). In this groundwater, two populations of *Methanoperedens* were identified, which could enable alternate roles in methane cycling.

Proteomic data supported episymbiosis of CPR bacteria with a host cell, as observed in cocultures of other CPR organisms with their respective hosts (62, 67, 68). Genomic data suggest a role for some CPR bacteria and DPANN archaea in sulfur cycling, possibly reflecting the metabolism of host cells (37, 62). In this groundwater sulfur cycling organisms belong to *Desulfobacterota* and *Nitrospirota*, but it is not yet known if DPANN archaea can have bacterial hosts (69). Proteins required for the formation of a cellulosome detected in *Cand*. Shapirobacterales indicate CPR bacteria also contribute to carbon cycling in this groundwater. Cellulosomes have been detected in *Cand*. Roizmanbacterium (70), a CPR organism also from the *Microgenomates* supergroup. *Cand*. Roizmanbacterium was similarly recovered from groundwater and encoded lactate dehydrogenase, suggesting a common role for groundwater *Microgenomates* in the production of labile organic carbon.

Our results demonstrate that methane and sulfur fuel anaerobic microbial metabolism in fracture water from the deep terrestrial subsurface. Sulfate and methane are abundant in many deep crystalline bedrock environments (19, 25, 71) which represent a significant fraction (∼20%) of the Earth’s surface (72). Syntrophy between methane and sulfur cycling organisms therefore has the potential to be widespread in SMTZs throughout the terrestrial biosphere, though the quantitative significance of the process remains to be determined. Carbon isotope values of DIC and fracture calcites at Olkiluoto indicate that methane oxidation is limited, with only one other sample showing evidence of a contribution from AOM (19). Thus, active methane oxidation may be a phenomenon that occurs locally in distinct locations throughout terrestrial SMTZs. The results further demonstrate the importance of activity measurements such as metaproteomics in genome-based studies that aim to uncover the roles of microorganisms in biogeochemical cycling. From only genome-based potential, our data would suggest that hydrogenotrophic sulfate reduction, which is common in other sulfidic Olkiluoto groundwaters (11), is the prominent metabolism with a limited contribution from ANME archaea, which were relatively low abundance in both the 16S rRNA gene amplicon and metagenomic datasets. With insight from metaproteomics, we propose that both anaerobic methane oxidation and sulfur disproportionation are important metabolisms in this deep oligotrophic groundwater.

## Materials and Methods

### Sample site and chemistry

Groundwater was collected from drillhole OL-KR13 on the island of Olkiluoto (61°14’30.7’’N, 21°28’49.3’’) from a fracture at 330.52–337.94 meters below sea level at ten specific time points for 9 months. The bedrock at Olkiluoto is predominantly granite and high-grade metamorphic rock (19). Hydrological models have shown that drillhole OL-KR13 intersects a fracture zone that connects to ONKALO, the underground nuclear waste repository that is currently under construction and that extends to ∼455 m depth (73). This connection can result in slow groundwater flow caused by drawdown towards open tunnels via the fracture zone.

Methods of groundwater chemical analysis have been described in detail previously (11). Briefly, anions (sulfate, thiosulfate, nitrate, nitrite) were measured by ion chromatography using a Dionex Integrion HPIC system with an IonPac AS11HC analytical column. Total and dissolved organic carbon were measured on a Shimadzu TOC-V. Organic acids (acetate, lactate, propionate, butyrate) and glucose were measured with an Agilent 1290 Infinity Liquid Chromatography System fitted with an Agilent Hi-Plex H column with RI detection. Alcohols (ethanol, methanol, propanol, and 2-butanol) and acetone were measured on a Varian CP-3800 gas chromatograph with 1-butanol as an internal standard. Dissolved gases (CH_4_ and H_2_) were measured on a Varian 450-GC with flame ionization (FID) and thermionic specific (TSD) detector. The ^12^C/^13^C ratio of methane (δ^13^C_CH4_ ‰ VPBD) was measured with a Picarro G2201-I Analyser. The ^12^C/^13^C ratio of dissolved inorganic carbon (DIC) (δ^13^C_DIC_ ‰ VPBD) and the ^32^S/^34^S isotope ratio of sulfate (δ^34^S_SO4_ ‰ VCDT) were measured at the Stable Isotope Laboratory, Faculty of Geosciences and Environment, Université de Lausanne, Switzerland. The C isotope composition was measured with a Thermo Finnigan Delta Plus XL IRMS equipped with a GasBench II for analyses of carbonates. The S isotope composition was measured with a He carrier gas and a Carlo Erba (CE 1100) elemental analyzer linked to a Thermo Fisher Delta mass spectrometer. Additional chemical measurements were conducted by Teollisuuden Voima Oyj (TVO), Finland, and methods are described in (74).

### Microbial biomass collection, DNA and protein extraction

To collect biomass for DNA and protein extraction, groundwater was pumped directly into a sterile chilled Nalgene filtration unit fitted with a 0.22 µm pore size Isopore polycarbonate membrane (Merck Millipore, Darmstadt, Germany). Approximately 10 L of groundwater was filtered each for metagenomic and metaproteomic sample and 1 L was filtered when samples were taken for 16S rRNA gene amplicon analysis only. Ultra-small cells in the 0.2 µm filtrate were collected by subsequent filtration through a 0.1 µm pore size polycarbonate filter. Filters collected for DNA were preserved in 750 µL LifeGuard Soil Preservation Solution (MoBio, Carlsbad, CA, United States) and stored at −20°C. Filters collected for protein were flash-frozen in a dry ice and ethanol mixture and stored at −80°C. DNA was extracted using a modified phenol-chloroform method (11, 75) and quantified with the dsDNA High Sensitivity Assay kit (Thermo Fisher Scientific) on a Qubit 3.0 Fluorometer. Protein was extracted at the Oak Ridge National Laboratory (Oak Ridge, TN, United States) using previously described methods (11, 76).

### DNA sequencing

Extracted DNA was used as a template for PCR amplification using primers 515F/806R that target the V4 region of the 16S rRNA gene (77). Amplicon libraries were sequenced on an Illumina MiSeq at either RTL Genomics, Lubbock, TX, USA, or the Lausanne Genomics Technologies Facility at Université de Lausanne (UNIL), Lausanne, Switzerland, with a 2 × 250 bp read configuration. Raw sequence reads were merged and quality-filtered with USEARCH v11 (78). Zero-radius operational taxonomic units (ZOTUs) were generated with UNOISE3 (79). Taxonomy was predicted with SINTAX (80) using a USEARCH compatible (81) SILVA 138 release (82). The resulting ZOTU table was analyzed and visualized with the R-package ampvis2 (83).

Seven metagenomes were sequenced at three facilities: The Joint Genome Institute (JGI), Walnut Creek, CA, USA; the Marine Biological Laboratory, Woods Hole, MA, USA, and; RTL Genomics, Lubbock, TX, USA. DNA was sequenced on an Illumina HiSeq 2500 (RTL and JGI) and Illumina NextSeq (MBL) with paired end 2 × 150 bp reads with further details in (11). All samples were pre-processed following the same procedure: low quality bases and adaptors were removed with cudadapt (84) and artificial PCR duplicates were screened using FastUniq (85) with default parameters.

### Metagenomic assembly, genome binning and single amplified genomes

After preprocessing, samples were individually assembled using MEGAHIT (86) with the -meta-sensitive option. Assembled contigs were binned using two methods: (1) assembled contigs were quantified across all samples using Kallisto (87) and binned using CONCOCT (88); (2) assembled contigs were quantified across all samples using bbmap (89) and binned using metabat2 (90). A non-redundant set of optimized bins was then selected with DAS Tool (91), with the default score threshold of 0.5. This resulted in a total of 238 bins from seven metagenomic datasets (June, July, September (×3), November (×2)). Taxonomy was assigned to bins with GTDB-Tk v1.3.0 reference data version r95 (92). The 238 bins were dereplicated with dRep (93) with parameters: minimum completeness 70%, maximum contamination 10%, primary cluster ANI 90%, secondary cluster ANI 99% (strain level). Bins belonging to *Patescibacteria* (50/238) were checked for completeness and contamination with the Candidate Phyla Radiation (CPR; *Patescibacteria*) custom 43 gene marker set in CheckM (94) and clustered into groups (ANI 99 %) with FastANI (95). In cases where the completeness of *Patescibacteria* bins improved past the threshold used for dereplication (70%), the bin with the greatest completeness and least contamination from each cluster was added to the collection of dereplicated bins from dRep. This resulted in a total of 76 dereplicated genome bins, hereafter referred to as metagenome-assembled genomes (MAGs).

Single amplified genomes (SAGs) were sequenced on an Illumina NextSeq. Raw reads were quality-filtered using BBTools (89) and assembled with SPAdes v3.13.0 (96). Contig ends were trimmed and discarded if the length was <2kb or the read coverage was less than 2. SAGs were clustered into groups (ANI 99%) with FastANI (95) and completeness and contamination was estimated with CheckM (94). Eight dereplicated SAGs were retained resulting in a total of 84 genomes (MAGs and SAGs) in our dataset. A phylogenomic tree was constructed with the 84 genomes from this study using GToTree (97) using Hidden Markov Models (HMMs) for 16 universal single copy genes (98). GTDB representative species from the same families as the groundwater MAGs and SAGs were recovered using gtt-get-accessions-from-GTDB and included in the tree. The tree was visualized with iToL (99).

### Metabolic predictions

Protein coding genes were predicted with Prodigal (100). METABOLIC v4.0 (101) was used to search amino acid sequences against a curated set of KOfam (102), TIGRfam (103), Pfam (104) and sulfur-related HMM profiles (37) corresponding to key marker genes for biogeochemical cycling. Genes putatively involved in sulfur disproportionation (YTD gene cluster consisting of a YEDE-like gene, *tusA*, a *dsrE*-like gene, and two conserved hypothetical proteins (39) were identified with custom HMMs (https://github.com/emma-bell/metabolism). For the generation of custom HMM profiles, reference gene sequences identified in (39) were aligned using MUSCLE (105) with default parameters. HMMs were built with hmmbuild in HMMER v3.3.2 (106). The *tusA* HMM was checked for CPxP conserved residues that stabilize the first helix (107). HMM profiles were searched against genomes using search-custom-markers workflow in metabolisHMM (108). Complexes putatively involved in metal oxide reduction by *Methanoperendaceae* (32) were identified by searching for CXXCH heme-binding motifs. Cellular location was predicted with PsortB v3.0 (109) and transmembrane helices were identified with TMHMM v2.0 (110). Putative function derived from CPR and DPANN protein sequences was predicted by searching conserved domains within protein coding sequences using CD-search (111) and performing similarity searches using blastp against the non-redundant protein sequences (nr) database (112).

### Metaproteomics

Digested peptides were analyzed by online two-dimensional liquid chromatography tandem mass spectrometry on an Orbitrap-Elite mass spectrometer (ThermoFisher Scientific). Mass spectra data were acquired in data dependent mode and the top ten most abundant parent ions were selected for further fragmentation by collision-induced dissociation at 35% energy level. Three technical replicates were analyzed per sample.

A protein database was constructed from predicted protein sequences from the seven metagenomic datasets. All collected MS/MS spectra were searched against the protein database with Myrimatch v2.2 (113). Peptides were identified and assembled into proteins using IDPicker v3.1 (114) with a minimum of two distinct peptides per protein and a false discovery rate (FDR) of <1% at the peptide level. Proteins were further clustered into protein groups post database searching if all proteins shared the same set of identified peptides.

## Data Availability

Supporting datasets are provided in the Supplementary Information (Datasets S1-S7). Sequence data are deposited in the National Centre for Biotechnology Information (NCBI) Sequence Read Archive (SRA). 16S rRNA gene amplicon data are deposited under the NCBI Bioproject PRJNA472445. BioSample accessions for MAGs and SAGs are provided in Dataset S2.

## Acknowledgments

We thank POSIVA OY for providing the funding and the logistical support to carry out the work, Maarit Yli-Kaila and Raila Viitala for support and assistance conducting fieldwork, Louise Balmer and Guillaume Sommer for assistance collecting samples, and Petteri Pitkänen for valuable discussions. Support for metagenomic sequencing was provided by The Census of Deep Life within the Deep Carbon Observatory and the U.S. Department of Energy Joint Genome Institute. The work conducted by the U.S. Department of Energy Joint Genome Institute, a DOE Office of Science User Facility, is supported under Contract No. DE-AC02-05CH11231. The metagenomic computations were performed on resources provided by SNIC through Uppsala Multidisciplinary Center for Advanced Computational Science (UPPMAX).

## Notes

### Competing Interest Statement

The authors have declared no competing interest.

## References

1. C. Magnabosco, et al., The biomass and biodiversity of the continental subsurface. Nat. Geosci. 11, 707–717 (2018).

2. T. M. Hoehler, B. B. Jørgensen, Microbial life under extreme energy limitation. Nat. Rev. Microbiol. 11, 83–94 (2013).

3. S. Braun, et al., Microbial turnover times in the deep seabed studied by amino acid racemization modelling. Sci. Rep. 7, 5680 (2017).

4. B. B. Jørgensen, Deep subseafloor microbial cells on physiological standby. Proc. Natl. Acad. Sci. 108, 18193LP–18194 (2011).

5. E. Trembath-Reichert, et al., Methyl-compound use and slow growth characterize microbial life in 2-km-deep subseafloor coal and shale beds. Proc. Natl. Acad. Sci. 114, E9206LP–E9215 (2017).

6. M. C. Y. Lau, et al., An oligotrophic deep-subsurface community dependent on syntrophy is dominated by sulfur-driven autotrophic denitrifiers. Proc. Natl. Acad. Sci. U. S. A. 113, E7927–E7936 (2016).

7. E. Bell, et al., Active sulfur cycling in the terrestrial deep subsurface. ISME J. (2020) https://doi.org/10.1038/s41396-020-0602-x.

8. M. Lopez-Fernandez, et al., Metatranscriptomes Reveal That All Three Domains of Life Are Active but Are Dominated by Bacteria in the Fennoscandian Crystalline Granitic Continental Deep Biosphere. MBio 9, e01792–18 (2018).

9. M. Mehrshad, et al., Energy efficiency and biological interactions define the core microbiome of deep oligotrophic groundwater. bioRxiv, 2020.05.24.111179 (2020).

10. P. Rajala, et al., Rapid Reactivation of Deep Subsurface Microbes in the Presence of C-1 Compounds. Microorganisms 3, 17–33 (2015).

11. E. Bell, et al., Biogeochemical cycling by a low-diversity microbial community in deep groundwater. Front. Microbiol. 9, 2129 (2018).

12. A. Vigneron, et al., Succession in the petroleum reservoir microbiome through an oil field production lifecycle. ISME J. 11, 2141–2154 (2017).

13. R. A. Daly, et al., Microbial metabolisms in a 2.5-km-deep ecosystem created by hydraulic fracturing in shales. Nat. Microbiol. 1, 16146 (2016).

14. S. Y. Park, Y. Liang, Biogenic methane production from coal: A review on recent research and development on microbially enhanced coalbed methane (MECBM). Fuel 166, 258–267 (2016).

15. A. Bagnoud, et al., Reconstructing a hydrogen-driven microbial metabolic network in Opalinus Clay rock in Nature Communications, (2016), p. 12770.

16. C. Hubert, “Microbial Ecology of Oil Reservoir Souring and its Control by Nitrate Injection” in Handbook of Hydrocarbon and Lipid Microbiology, (Springer Berlin Heidelberg, 2010), pp. 2753–2766.

17. S. Lahme, J. Mand, J. Longwell, R. Smith, D. Enning, Severe Corrosion of Carbon Steel in Oil Field Produced Water Can Be Linked to Methanogenic Archaea Containing a Special Type of [NiFe] Hydrogenase. Appl. Environ. Microbiol. 87, e01819–20 (2021).

18. L. M. Gieg, T. R. Jack, J. M. Foght, Biological souring and mitigation in oil reservoirs. Appl. Microbiol. Biotechnol. 92, 263–282 (2011).

19. Posiva, Olkiluoto Site description, Posiva report 2011-02 (2013).

20. K. Knittel, A. Boetius, Anaerobic Oxidation of Methane: Progress with an Unknown Process. Annu. Rev. Microbiol. 63, 311–334 (2009).

21. M. Egger, N. Riedinger, J. M. Mogollón, B. B. Jørgensen, Global diffusive fluxes of methane in marine sediments. Nat. Geosci. 11, 421–425 (2018).

22. A. J. Wallenius, P. Dalcin Martins, C. P. Slomp, M. S. M. Jetten, Anthropogenic and Environmental Constraints on the Microbial Methane Cycle in Coastal Sediments. Front. Microbiol. 12, 293 (2021).

23. M. Bomberg, M. Nyyssönen, P. Pitkänen, A. Lehtinen, M. Itävaara, Active Microbial Communities Inhabit Sulphate-Methane Interphase in Deep Bedrock Fracture Fluids in Olkiluoto, Finland. Biomed Res. Int. 2015, 1–17 (2015).

24. M. Bomberg, T. Lamminmäki, M. Itävaara, Microbial communities and their predicted metabolic characteristics in deep fracture groundwaters of the crystalline bedrock at Olkiluoto, Finland. Biogeosciences 13 (2016).

25. K. Ino, et al., Deep microbial life in high-quality granitic groundwater from geochemically and geographically distinct underground boreholes. Environ. Microbiol. Rep. 8 (2016).

26. K. Ino, et al., Ecological and genomic profiling of anaerobic methane-oxidizing archaea in a deep granitic environment. ISME J. 12, 31–47 (2018).

27. A. W. Hernsdorf, et al., Potential for microbial H2 and metal transformations associated with novel bacteria and archaea in deep terrestrial subsurface sediments. ISME J. 11, 1915–1929 (2017).

28. H. Drake, et al., Extreme 13C depletion of carbonates formed during oxidation of biogenic methane in fractured granite. Nat. Commun. 6, 7020 (2015).

29. A. O. Leu, et al., Lateral Gene Transfer Drives Metabolic Flexibility in the Anaerobic Methane-Oxidizing Archaeal Family *Methanoperedenaceae*; MBio 11, e01325–20 (2020).

30. K. F. Ettwig, et al., Archaea catalyze iron-dependent anaerobic oxidation of methane. Proc. Natl. Acad. Sci. 113, 12792–12796 (2016).

31. M. F. Haroon, et al., Anaerobic oxidation of methane coupled to nitrate reduction in a novel archaeal lineage. Nature 500, 567–570 (2013).

32. A. O. Leu, et al., Anaerobic methane oxidation coupled to manganese reduction by members of the Methanoperedenaceae. ISME J. (2020) https://doi.org/10.1038/s41396-020-0590-x.

33. H. S. Weber, K. S. Habicht, B. Thamdrup, Anaerobic methanotrophic archaea of the ANME-2d cluster are active in a low-sulfate, iron-rich freshwater sediment. Front. Microbiol. 8, 619 (2017).

34. G. Su, et al., Manganese/iron-supported sulfate-dependent anaerobic oxidation of methane by archaea in lake sediments. Limnol. Oceanogr. 65, 863–875 (2020).

35. K. Pedersen, Metabolic activity of subterranean microbial communities in deep granitic groundwater supplemented with methane and H 2. ISME J. 7, 839–849 (2013).

36. C. Dahl, C. Friedrich, A. Kletzin, “Sulfur Oxidation in Prokaryotes” in Encyclopedia of Life Sciences, (John Wiley & Sons, Ltd, 2008) https://doi.org/10.1002/9780470015902.a0021155 (January 21, 2019).

37. K. Anantharaman, et al., Expanded diversity of microbial groups that shape the dissimilatory sulfur cycle. ISME J. 12, 1715–1728 (2018).

38. M. Motamedi, K. Pedersen, Note: Desulfovibrio aespoeensis sp. nov., a mesophilic sulfate-reducing bacterium from deep groundwater at aspo hard rock laboratory, Sweden. Int. J. Syst. Bacteriol. 48, 311–315 (1998).

39. K. Umezawa, H. Kojima, Y. Kato, M. Fukui, Disproportionation of inorganic sulfur compounds by a novel autotrophic bacterium belonging to Nitrospirota. Syst. Appl. Microbiol. 43, 126110 (2020).

40. K. Finster, Microbiological disproportionation of inorganic sulfur compounds. J. Sulfur Chem. 29, 281–292 (2008).

41. C. Thorup, A. Schramm, A. J. Findlay, K. W. Finster, L. Schreiber, Disguised as a Sulfate Reducer: Growth of the Deltaproteobacterium Desulfurivibrio alkaliphilus by Sulfide Oxidation with Nitrate. MBio 8, e00671–17 (2017).

42. A. I. Slobodkin, G. B. Slobodkina, Diversity of Sulfur-Disproportionating Microorganisms. Microbiol. (Russian Fed. 88, 509–522 (2019).

43. K. W. Finster, et al., Complete genome sequence of Desulfocapsa sulfexigens, a marine deltaproteobacterium specialized in disproportionating inorganic sulfur compounds. Stand. Genomic Sci. 8, 58–68 (2013).

44. M. Jormakka, et al., Molecular mechanism of energy conservation in polysulfide respiration. Nat. Struct. Mol. Biol. 15, 730–737 (2008).

45. S. Duval, A.-L. Ducluzeau, W. Nitschke, B. Schoepp-Cothenet, Enzyme phylogenies as markers for the oxidation state of the environment: the case of respiratory arsenate reductase and related enzymes. BMC Evol. Biol. 8, 206 (2008).

46. J. E. Butler, N. D. Young, D. R. Lovley, Evolution of electron transfer out of the cell: comparative genomics of six Geobacter genomes. BMC Genomics 11, 40 (2010).

47. C. Cai, et al., A methanotrophic archaeon couples anaerobic oxidation of methane to Fe(III) reduction. ISME J. 12, 1929–1939 (2018).

48. S. E. McGlynn, G. L. Chadwick, C. P. Kempes, V. J. Orphan, Single cell activity reveals direct electron transfer in methanotrophic consortia. Nature 526, 531–535 (2015).

49. V. Krukenberg, et al., Gene expression and ultrastructure of meso-and thermophilic methanotrophic consortia. Environ. Microbiol. 20, 1651–1666 (2018).

50. D. Søndergaard, C. N. S. Pedersen, C. Greening, HydDB: A web tool for hydrogenase classification and analysis. Sci. Rep. 6, 34212 (2016).

51. P. Pedroni, et al., Characterization of the locus encoding the [Ni-Fe] sulfhydrogenase from the archaeon Pyrococcus furiosus: evidence for a relationship to bacterial sulfite reductases. Microbiology 141, 449–458 (1995).

52. K. Ma, R. N. Schicho, R. M. Kelly, M. W. Adams, Hydrogenase of the hyperthermophile Pyrococcus furiosus is an elemental sulfur reductase or sulfhydrogenase: evidence for a sulfur-reducing hydrogenase ancestor. Proc. Natl. Acad. Sci. 90, 5341LP–5344 (1993).

53. B. Luef, et al., Diverse uncultivated ultra-small bacterial cells in groundwater. Nat. Commun. 6, 6372 (2015).

54. C. J. Castelle, et al., Biosynthetic capacity, metabolic variety and unusual biology in the CPR and DPANN radiations. Nat. Rev. Microbiol. 16, 629–645 (2018).

55. K. C. Wrighton, et al., Fermentation, hydrogen, and sulfur metabolism in multiple uncultivated bacterial phyla. Science 337, 1661–5 (2012).

56. C. J. Castelle, et al., Genomic Expansion of Domain Archaea Highlights Roles for Organisms from New Phyla in Anaerobic Carbon Cycling. Curr. Biol. 25, 690–701 (2015).

57. R. E. Danczak, et al., Members of the Candidate Phyla Radiation are functionally differentiated by carbon-and nitrogen-cycling capabilities. Microbiome 5, 112 (2017).

58. L. Tišáková, B. Vidová, J. Farkašovská, A. Godány, Bacteriophage endolysin Lyt μ1/6: characterization of the C-terminal binding domain. FEMS Microbiol. Lett. 350, 199–208 (2014).

59. B. Maier, G. C. L. Wong, How Bacteria Use Type IV Pili Machinery on Surfaces. Trends Microbiol. 23, 775–788 (2015).

60. K. C. Wrighton, et al., Metabolic interdependencies between phylogenetically novel fermenters and respiratory organisms in an unconfined aquifer. ISME J. 8, 1452–1463 (2014).

61. K. C. Wrighton, et al., RubisCO of a nucleoside pathway known from Archaea is found in diverse uncultivated phyla in bacteria. ISME J. 10, 2702–2714 (2016).

62. C. He, et al., Genome-resolved metagenomics reveals site-specific diversity of episymbiotic CPR bacteria and DPANN archaea in groundwater ecosystems. Nat. Microbiol. (2021) https://doi.org/10.1038/s41564-020-00840-5.

63. L. Johansson, J. Stahlén, T. Taborowski, K. Pedersen, “Microbial Release of Iron from Olkiluoto Rock Minerals” (Posiva Oy, 2019).

64. J. Milucka, et al., Zero-valent sulphur is a key intermediate in marine methane oxidation. Nature 491, 541–546 (2012).

65. S.-C. Chen, et al., Anaerobic oxidation of ethane by archaea from a marine hydrocarbon seep. Nature 568, 108–111 (2019).

66. R. Kevorkian, S. Callahan, R. Winstead, K. G. Lloyd, ANME-1 archaea drive methane accumulation and removal in estuarine sediments. bioRxiv, 2020.02.24.963215 (2020).

67. X. He, et al., Cultivation of a human-associated TM7 phylotype reveals a reduced genome and epibiotic parasitic lifestyle. Proc. Natl. Acad. Sci. 112, 244LP–249 (2015).

68. K. L. Cross, et al., Targeted isolation and cultivation of uncultivated bacteria by reverse genomics. Nat. Biotechnol. 37, 1314–1321 (2019).

69. C. J. Castelle, et al., Protein family content uncovers lineage relationships and bacterial pathway maintenance mechanisms in DPANN archaea. bioRxiv, 2021.01.12.426361 (2021).

70. P. Geesink, et al., Genome-inferred spatio-temporal resolution of an uncultivated Roizmanbacterium reveals its ecological preferences in groundwater. Environ. Microbiol. 22, 726–737 (2020).

71. R. Kietäväinen, L. Purkamo, The origin, source, and cycling of methane in deep crystalline rock biosphere. Front. Microbiol. 6, 725 (2015).

72. S. Chandra, E. Auken, P. K. Maurya, S. Ahmed, S. K. Verma, Large Scale Mapping of Fractures and Groundwater Pathways in Crystalline Hardrock By AEM. Sci. Rep. 9, 398 (2019).

73. T. Vaittinen, H. Ahokas, J. Nummela, S. Paulamäki, “Hydrogeological Structure Model of the Olkiluoto Site – Update in 2010, Posiva Report 2011-65” (2011).

74. P. Tuomi, et al., “Conceptual Model of Microbial Effects on Hydrogeochemical Conditions at the Olkiluoto Site” in Posiva Report 2020-03, (Posiva Oy, 2020), p. 278.

75. A. Bagnoud, O. Leupin, B. Schwyn, R. Bernier-Latmani, Rates of microbial hydrogen oxidation and sulfate reduction in Opalinus Clay rock. Appl. Geochemistry 72, 42–50 (2016).

76. K. Chourey, et al., Environmental proteomics reveals early microbial community responses to biostimulation at a uranium-and nitrate-contaminated site. Proteomics 13, n/a-n/a (2013).

77. J. G. Caporaso, et al., Global patterns of 16S rRNA diversity at a depth of millions of sequences per sample. Proc. Natl. Acad. Sci. 108, 4516–4522 (2011).

78. R. C. Edgar, Search and clustering orders of magnitude faster than BLAST. Bioinformatics 26, 2460–2461 (2010).

79. R. C. Edgar, “UNOISE2: improved error-correction for Illumina 16S and ITS amplicon sequencing” (Cold Spring Harbor Laboratory, 2016) https://doi.org/10.1101/081257 (October 3, 2017).

80. R. Edgar, SINTAX: a simple non-Bayesian taxonomy classifier for 16S and ITS sequences. bioRxiv, 074161 (2016).

81. M. Lee, making-usearch-compatible-tax-file-from-dada2-silva-v138-format.sh (2020) https://doi.org/10.6084/m9.figshare.12226943.v1.

82. C. Quast, et al., The SILVA ribosomal RNA gene database project: Improved data processing and web-based tools. Nucleic Acids Res. 41, D590–D596 (2013).

83. K. S. Andersen, R. H. Kirkegaard, S. M. Karst, M. Albertsen, ampvis2: an R package to analyse and visualise 16S rRNA amplicon data. bioRxiv, 299537 (2018).

84. M. Martin, Cutadapt removes adapter sequences from high-throughput sequencing reads. EMBnet.journal; Vol 17, No 1 Next Gener. Seq. Data Anal. (2011) https://doi.org/10.14806/ej.17.1.200.

85. H. Xu, et al., FastUniq: A Fast De Novo Duplicates Removal Tool for Paired Short Reads. PLoS One 7, e52249 (2012).

86. D. Li, C. M. Liu, R. Luo, K. Sadakane, T. W. Lam, MEGAHIT: An ultra-fast single-node solution for large and complex metagenomics assembly via succinct de Bruijn graph. Bioinformatics 31, 1674–1676 (2015).

87. N. L. Bray, H. Pimentel, P. Melsted, L. Pachter, Near-optimal probabilistic RNA-seq quantification. Nat. Biotechnol. 34, 525–527 (2016).

88. J. Alneberg, et al., Binning metagenomic contigs by coverage and composition. Nat. Methods 11, 1144–1146 (2014).

89. B. Bushnell, J. Rood, E. Singer, BBTools Software Package. PLoS One 12, e0185056 (2017).

90. D. D. Kang, et al., MetaBAT 2: an adaptive binning algorithm for robust and efficient genome reconstruction from metagenome assemblies. PeerJ 7, e7359–e7359 (2019).

91. C. M. K. K. Sieber, et al., Recovery of genomes from metagenomes via a dereplication, aggregation and scoring strategy. Nat. Microbiol. 3, 836–843 (2018).

92. P.-A. Chaumeil, A. J. Mussig, P. Hugenholtz, D. H. Parks, GTDB-Tk: a toolkit to classify genomes with the Genome Taxonomy Database. Bioinformatics (2019) https://doi.org/10.1093/bioinformatics/btz848 (January 9, 2020).

93. M. R. Olm, C. T. Brown, B. Brooks, J. F. Banfield, dRep: a tool for fast and accurate genomic comparisons that enables improved genome recovery from metagenomes through de-replication. ISME J. 11, 2864–2868 (2017).

94. D. H. Parks, M. Imelfort, C. T. Skennerton, P. Hugenholtz, G. W. Tyson, CheckM: Assessing the quality of microbial genomes recovered from isolates, single cells, and metagenomes. Genome Res. 25, 1043–1055 (2015).

95. C. Jain, L. M. Rodriguez-R, A. M. Phillippy, K. T. Konstantinidis, S. Aluru, High throughput ANI analysis of 90K prokaryotic genomes reveals clear species boundaries. Nat. Commun. 9, 5114 (2018).

96. A. Bankevich, et al., SPAdes: a new genome assembly algorithm and its applications to single-cell sequencing. J. Comput. Biol. 19, 455–77 (2012).

97. M. D. Lee, GToTree: a user-friendly workflow for phylogenomics. Bioinformatics (2019) https://doi.org/10.1093/bioinformatics/btz188 (July 18, 2019).

98. L. A. Hug, et al., Critical biogeochemical functions in the subsurface are associated with bacteria from new phyla and little studied lineages. Environ. Microbiol. 18, 159–173 (2016).

99. I. Letunic, P. Bork, Interactive Tree Of Life (iTOL) v5: an online tool for phylogenetic tree display and annotation. Nucleic Acids Res. 49, W293–W296 (2021).

100. D. Hyatt, et al., Prodigal: prokaryotic gene recognition and translation initiation site identification. BMC Bioinformatics 11, 119 (2010).

101. Z. Zhou, et al., METABOLIC: High-throughput profiling of microbial genomes for functional traits, biogeochemistry, and community-scale metabolic networks. bioRxiv, 761643 (2020).

102. T. Aramaki, et al., KofamKOALA: KEGG Ortholog assignment based on profile HMM and adaptive score threshold. Bioinformatics 36, 2251–2252 (2020).

103. J. D. Selengut, et al., TIGRFAMs and Genome Properties: tools for the assignment of molecular function and biological process in prokaryotic genomes. Nucleic Acids Res. 35, D260–D264 (2007).

104. R. D. Finn, et al., Pfam: the protein families database. Nucleic Acids Res. 42, D222–D230 (2014).

105. R. C. Edgar, MUSCLE: multiple sequence alignment with high accuracy and high throughput. Nucleic Acids Res. 32, 1792–1797 (2004).

106. S. R. Eddy, Accelerated Profile HMM Searches. PLoS Comput. Biol. 7, e1002195–e1002195 (2011).

107. E. Katoh, et al., High precision NMR structure of YhhP, a novel Escherichia coli protein implicated in cell division11 Edited by M. F. Summers. J. Mol. Biol. 304, 219–229 (2000).

108. E. A. McDaniel, K. Anantharaman, K. D. McMahon, metabolisHMM: Phylogenomic analysis for exploration of microbial phylogenies and metabolic pathways. bioRxiv, 2019.12.20.884627 (2019).

109. N. Y. Yu, et al., PSORTb 3.0: improved protein subcellular localization prediction with refined localization subcategories and predictive capabilities for all prokaryotes. Bioinformatics 26, 1608–1615 (2010).

110. A. Krogh, B. Larsson, G. von Heijne, E. L. L. Sonnhammer, Predicting transmembrane protein topology with a hidden markov model: application to complete genomes11Edited by F. Cohen. J. Mol. Biol. 305, 567–580 (2001).

111. S. Lu, et al., CDD/SPARCLE: the conserved domain database in 2020. Nucleic Acids Res. 48, D265–D268 (2020).

112. S. F. Altschul, W. Gish, W. Miller, E. W. Myers, D. J. Lipman, Basic local alignment search tool. J Mol Biol 215, 403–410 (1990).

113. D. L. Tabb, C. G. Fernando, M. C. Chambers, MyriMatch: Highly accurate tandem mass spectral peptide identification by multivariate hypergeometric analysis. J. Proteome Res. 6, 654–661 (2007).

114. Z. Q. Ma, et al., IDPicker 2.0: Improved protein assembly with high discrimination peptide identification filtering. J. Proteome Res. 8, 3872–3881 (2009).

